# kGWASflow: a modular, flexible, and reproducible Snakemake workflow for k-mers-based GWAS

**DOI:** 10.1101/2023.07.10.548365

**Authors:** Adnan Kivanc Corut, Jason G. Wallace

## Abstract

Genome-wide association studies (GWAS) have been widely used to identify genetic variation associated with complex traits. Despite its success and popularity, the traditional GWAS approach comes with a variety of limitations. For this reason, newer methods for GWAS have been developed, including the use of pan-genomes instead of a reference genome and the utilization of markers beyond single-nucleotide polymorphisms, such as structural variations and k-mers. The k-mers based GWAS approach has especially gained attention from researchers in recent years. However, these new methodologies can be complicated and challenging to implement. Here we present kGWASflow, a modular, user-friendly, and scalable workflow to perform GWAS using k-mers. We adopted an existing kmersGWAS method into an easier and more accessible workflow using management tools like Snakemake and Conda and eliminated the challenges caused by missing dependencies and version conflicts. kGWASflow increases the reproducibility of the kmersGWAS method by automating each step with Snakemake and using containerization tools like Docker. The workflow encompasses supplemental components such as quality control, read-trimming procedures, and generating summary statistics. kGWASflow also offers post-GWAS analysis options to identify the genomic location and context of trait-associated k-mers. kGWASflow can be applied to any organism and requires minimal programming skills. kGWASflow is freely available on GitHub (https://github.com/akcorut/kGWASflow) and Bioconda (https://anaconda.org/bioconda/kgwasflow).

## Introduction

Identifying genotype-phenotype associations is fundamental to understanding the genetic architecture of complex traits. Genomewide association studies (GWAS) have been the method of choice to detect associations between genetic variants and phenotypes for over 15 years (Visscher et al. 2017). Through GWAS, thousands of traits have been surveyed, and numerous statistically significant associations have been reported (Uffelmann et al. 2021; MacArthur *et al*. 2017). These findings resulted in a better understanding of complex human traits and diseases (Cano-Gamez and Trynka 2020), helped improve plant breeding (Tibbs Cortes et al. 2021) and animal health (Tian et al. 2020), and have otherwise significantly impacted our understanding of genetics.

The classical GWAS approach uses genome-wide single nucleotide polymorphisms (SNPs) as the genotype data. During a standard GWAS, SNP markers are tested for statistically significant association with the phenotypic trait using statistical models. GWAS utilizes linkage disequilibrium (LD) information between markers and causal variants to identify trait-associated loci. However, despite its power and success, this method comes with various limitations (Tam et al. 2019; Gupta et al. 2019; Sun *et al*. 2021). It has been previously shown that GWAS often fails to pinpoint causal variants due to linkage disequilibrium (Boyle et al. 2017; Faye et al. 2013; LaPierre et al. 2021). Additionally, these studies primarily rely on SNPs as markers of genetic variants and often ignore other variants, such as structural variations, and therefore can sometimes explain only a fraction of heritability, particularly in cases involving highly complex traits (Manolio *et al*. 2009; Nolte *et al*. 2017). GWA studies can also identify spurious associations (Sul *et al*. 2018) and fail to capture associations caused by rare variants (Wray *et al*. 2011; Young 2019).

The quality of GWAS also depends on the availability and the quality of reference genomes. Traditional GWAS relies on mapping sequencing reads to a reference genome and then calling variants. This mapping step can potentially cause biases during variant calling because reference genomes are frequently incomplete and may not represent the full spectrum of genetic variation within a population. In addition, the misalignments can result in incorrect variant calling, especially in complex genomes and/or around repetitive regions.

Due to these limitations, newer GWAS methods have been developed (Gupta 2021; Coletta *et al*. 2021). These newer methods include but are not limited to using pan-genomes instead of a single reference genome (Manuweera *et al*. 2019; Song *et al*. 2020; Zhou *et al*. 2022) and the usage of new markers beyond SNPs, such as structural variations (Prinsen *et al*. 2017; Zhou *et al*. 2018; Yang *et al*. 2019; Li *et al*. 2020; Qin *et al*. 2021; Göktay *et al*. 2021; Wei *et al*. 2021) and *k*-mers (Rahman *et al*. 2018; Mehrab *et al*. 2021; Voichek and Weigel 2020; He *et al*. 2021; Lemane *et al*. 2022). Using structural variations may capture some of the missing heritability (Theunissen *et al*. 2020; Génin 2020; Zhou *et al*. 2022). Furthermore, utilizing k-mers as genetic markers offers significant advantages compared to the traditional SNP-based approach. *k*-mers, substrings of length *k* in sequencing reads, can mark a broader range of genomic variants, including structural variations. k-mers-based GWAS also allows a reference-free association mapping and can identify trait-associated markers even in regions missing in the reference genome (Gupta 2021).

Voichek et al. (2020) recently developed a k-mers-based GWAS approach for both categorical and quantitative phenotypes (Voichek and Weigel 2020). In this approach, the authors use the presence or absence of k-mers in sequencing reads as genotypic variants and then apply traditional GWAS methods. Even though the k-mers-based GWAS method recently gained increased attention (Colque-Little et al. 2021; Tripodi et al. 2021; Schulthess *et al*. 2022; Kale et al. 2022; Onetto et al. 2022; Li et al. 2022), it has not reached its potential due to the difficulties in implementing it. These difficulties include the need for bioinformatics expertise and lack of user-friendly tools. Underlying software and library dependencies also have the potential to cause reproducibility issues. Moreover, the existing implementation of this method lacks the essential downstream analyses necessary for interpreting results from the k-mers-based GWAS approach.

In an effort to tackle these challenges, here we present kGWASflow, a modular, user-friendly, and scalable workflow to perform k-mers-based GWAS. kGWASflow adapts the approach by Voichek et al. (2020), creating a more user-friendly workflow while addressing software dependency and version conflict issues. Our workflow is highly deployable in high-performance and cloud computing environments. It also enhances the interpretation of k-mer-based GWAS findings by providing supplementary downstream analysis options. kGWASflow boasts extensive customization, providing users with a variety of options tailored to their specific requirements.

## Methods

### Overview

The overall workflow of kGWASflow is described in Figure 1. With the default settings, kGWASflow comprises three main phases: preprocessing, k-mers-based GWAS (Voichek and Weigel 2020), and post-GWAS analysis. In short, the pre-processing phase conducts quality control analysis, offers optional read trimming, and organizes the input files for downstream analysis. The workflow’s second and main phase performs k-mers-based GWAS by implementing the kmersGWAS method from Voichek et al. (2020) (Voichek and Weigel 2020). The post-GWAS phase generates results tables and provides multiple options to identify genomic locations and context of trait-associated *k*-mers. Lastly, kGWASflow generates an HTML report that includes QC and summary statistics, diagnostic plots, and kmersGWAS results. kGWASflow is highly customizable, as multiple steps of the workflow are optional and can be easily deactivated. kGWASflow is written in Snakemake (Mölder et al. 2021), a commonly used Python-based workflow engine.

**Figure 1.**
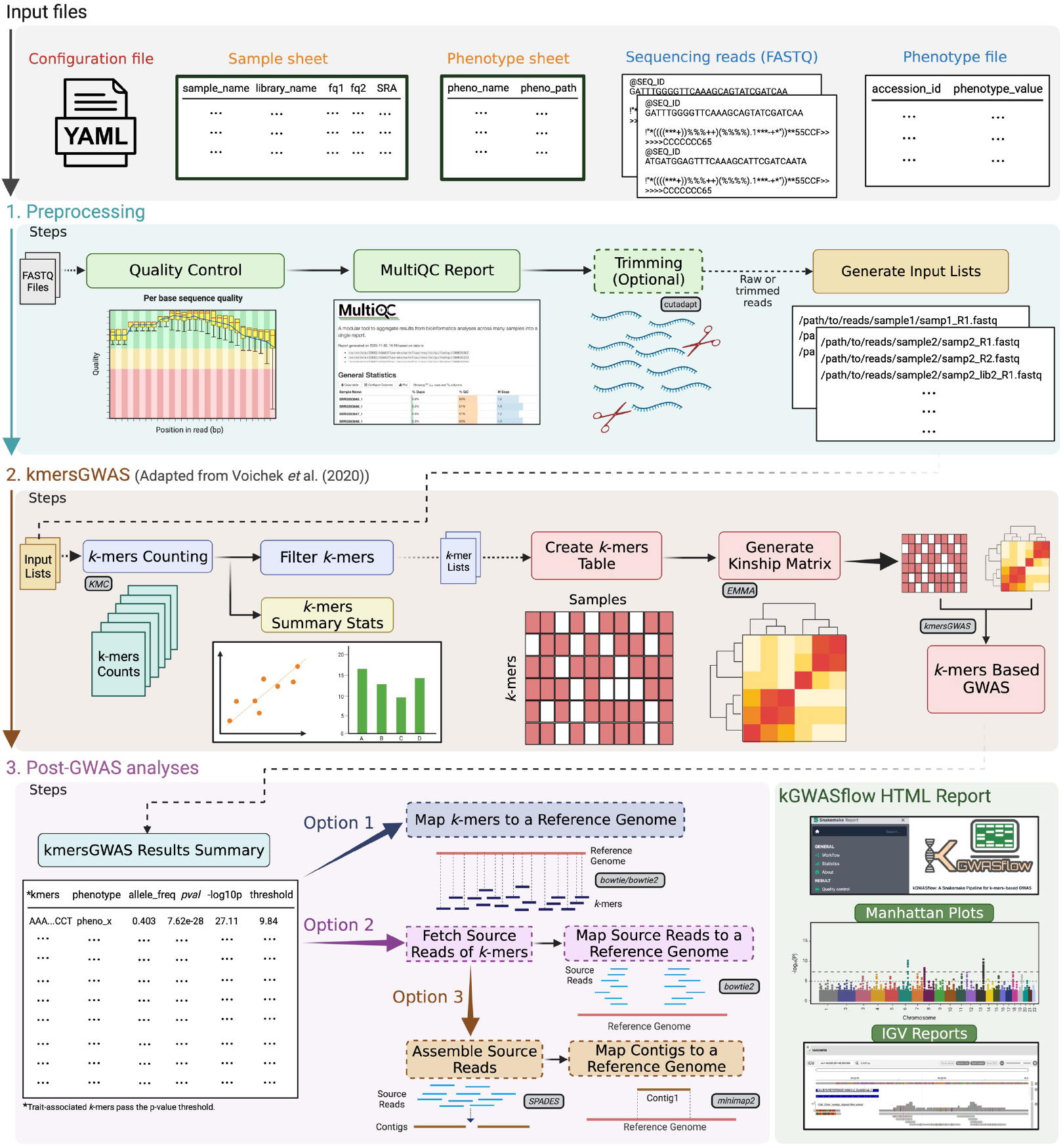
Overview of the kGWASflow workflow. The default kGWASflow workflow consists of three main phases: Preprocessing (highlighted in blue), k-mers-based GWAS (highlighted in brown), and post-GWAS analyses (highlighted in pink). The configuration and input files required by the workflow are highlighted in grey (top). The final outputs and report of the workflow are outlined in the bottom right corner (highlighted in green). Boxes outlined in black and filled in grey denote the publicly available tools employed in the workflow. The workflow steps are customizable, with multiple optional steps, such as read trimming and post-GWAS analysis options (options 1, 2, and 3). These optional steps are in dashed boxes.

Snakemake allows the workflow to be highly modular, scalable, and reproducible. Utilizing Snakemake, the workflow is constructed through a set of rules that link input sets with their corresponding outputs. Snakemake establishes the execution order by discerning the optimal combination of rules necessary to produce the target output while ensuring that each rule is triggered only upon the availability of its input files. Snakemake enhances the parallelization of independent jobs, considering computational resources and the thread requirements for each job’s execution. This feature enables automatic scalability, allowing users to effectively run the workflow on local machines and high-performance and cloud-based computing platforms. Another aspect of Snakemake is that it recognizes which output files are missing if the execution of the pipeline is halted due to an error and allows users to recover the execution of failed jobs and continue after the issues are resolved. By combining Snakemake with the Conda software manager, kGWASflow automatically installs all software and library dependencies for each rule separately in a Conda environment. In doing so, kGWASflow effectively addresses potential complications arising from software dependency conflicts across various stages of the pipeline. kGWASflow also takes advantage of Conda for the initial deployment of the workflow environment. The latest version of the workflow and its dependencies can be easily installed and activated by running the command

~~~
     conda create -c bioconda \
                    --name kgwasflow kgwasflow
     conda activate kgwasflow
~~~

### Input

The workflow has two main inputs: paired-end FASTQ files and single or multiple phenotype files. FASTQ files contain the sequencing reads of each individual/sample, and each individual/sample can have multiple units of FASTQ files obtained from different sequencing runs of the same sample. A separate phenotype file for each phenotype tested needs to be provided by the user. This phenotype file consists of two columns where the first column represents the name of the individual/sample and the second column represents the phenotypic value of that corresponding sample, as described in Voichek et al. (2020) (Voichek and Weigel 2020).

By configuring a single samples sheet (samples.tsv, Supplementary Data S5), users can easily supply all necessary sample information for the pipeline, including sample names, FASTQ file paths, or SRA accessions. Users have the option to either specify the local file path for each FASTQ file or supply SRA accessions for each sample, considering individual sequencing runs for the same sample separately. When provided with only SRA accessions, the pipeline automatically retrieves the relevant FASTQ files for each sample and its associated sequencing run using fasterq-dump. Phenotype file information can be provided by using the phenotype sheet (phenos.tsv, Supplementary Data S6). In the phenotype sheet, users specify the phenotype name in the first column and the file path of the corresponding phenotype file in the second column.

### Workflow configuration and execution

KGWASflow streamlines the workflow configuration process and provides users with easy customization. Once the workflow is installed, a new kGWASflow working directory with default configuration files can be initialized with a single command.

~~~
    kgwasflow init --work-dir working_dir/
~~~

This command will generate the configuration file ‘config.yaml’ (Supplementary Data S3) alongside tab-separated sample and phenotype sheets (explained in the Input section) inside of the config directory. The command will also generate the ‘test’ directory containing all the essential files required for executing a test run of kGWASflow. An example directory structure initialized by init command can be found in Figure 2.

**Figure 2.**
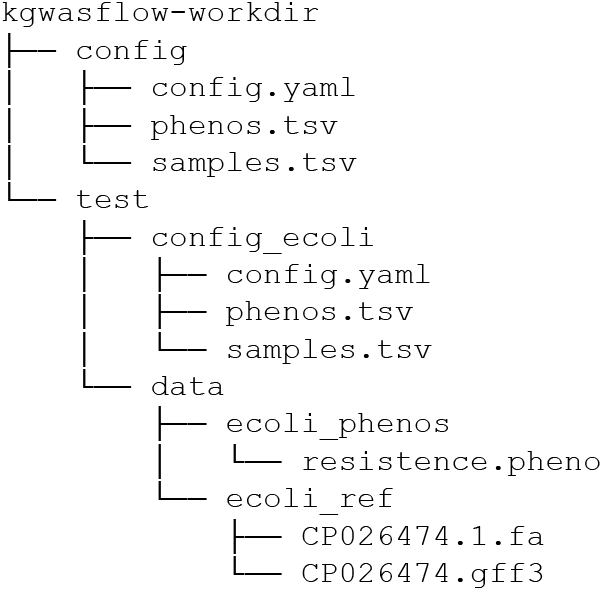
Example working directory structure generated by kG-WASflow initialization command: kgwasflow init. This working directory contains the default kGWASflow configuration files and the test/ directory with all the essential files required for a test workflow run.

To configure the workflow, the user simply needs to modify the ‘config.yaml’ configuration file, which is composed of three main sections: input information, workflow settings, and tool parameters. In the first section, the user defines the paths for the sample and phenotype sheets and provides details regarding the reference genome, if required. The workflow settings section allows users to control optional pipeline steps, such as read trimming, by changing them to ‘True’ or ‘False’ to activate or deactivate. In this section, users can also specify their preferred post-GWAS analysis options (Figure 1). Finally, in the tool parameters section, the users can define the parameters/settings for each tool used in the workflow.

After the configuration, kGWASfow can be run with a single command

~~~
    kgwasflow run --threads 16 \
                   --work-dir working_dir
~~~

This command first initiates the installation of all necessary software and library dependencies for the workflow via Conda before executing the workflow steps. The execution order is determined by Snakemake, according to the user-defined configuration file and the dependencies between the workflow steps.

Alternatively, the workflow can be implemented by cloning the GitHub repository and executing snakemake command after the configuration of the workflow. The configuration can be done by manually configuring the configuration files within the cloned repository. A detailed guide on how to install, configure, and use the pipeline via Conda or GitHub can be found on the kGWASflow wiki page (https://github.com/akcorut/kGWASflow/wiki).

### Workflow steps

#### Pre-processing

The pre-processing phase of kGWASflow starts with the quality control (QC) analysis of raw sequencing reads. kGWASflow first uses FastQC (Andrews 2010) to generate basic QC metrics, such as quality scores, GC content, duplication levels, and adapter content, for each FASTQ file. The pipeline then uses MultiQC (Ewels et al. 2016) to summarize and visualize each FastQC report in a single HTML file. If the trimming setting is activated, kGWASflow performs read trimming for each FASTQ file using cutadapt (Martin 2011). Users can define the parameters for read trimming in the configuration file. If the trimming setting is not activated, the pipeline skips this step and uses the raw reads for the rest of the workflow. During the final stage of the pre-processing phase, kGWASflow sorts the FASTQ files (trimmed or raw) into individual folders based on the sample or individual name, with one folder designated for each. Subsequently, within each folder, a text file is generated, listing the file paths of all FASTQ files associated with that particular sample. This last step is required to run *k*-mers-based GWAS.

#### k-mers based GWAS

Once the pre-processing phase is complete, kGWASflow starts the *k*-mers counting step. KMC (Kokot *et al*. 2017) is used to count *k*-mers from sequencing reads (trimmed or raw) from each individual/sample. Users can specify the desired *k*-mer length and the read count threshold parameters in the configuration file. Initially, KMC is run in default mode to count canonical *k*-mers, followed by a second run using the “-b” option to count non-canonical *k*-mers. Canonization in this context means that KMC, in its default mode, considers a *k*-mer and its reverse complement as equivalent and assigns the combined count of the two to the alphabetically smaller *k*-mer (Deorowicz *et al*. 2015). Conversely, in non-canonical counting mode (“-b” option), the k-mer and its reverse complement are counted separately. Next, the “kmers_add_strand_information” function from the kmersGWAS library is used to combine the output of two KMC runs. This generates a single list of k-mers per sample, along with their strand information. Subsequently, *k*-mer lists from each sample are merged into a single binary file (kmer_to_use) and then filtered using the “list_kmers_found_in_multiple_samples” feature of the kmersGWAS library. As described in Voichek et al. (2020), the two-step filtering criteria are applied as follows: First, *k*-mers are filtered if they are not present in at least N individuals. Second, *k*-mers are filtered based on their canonical and non-canonical counts. A *k*-mer is dropped if it is not found in both canonical and non-canonical forms in at least X percent of the individuals/samples in which it appeared (Voichek and Weigel 2020). Users can easily modify filtering criteria within the configuration file. Upon completing the *k*-mer counting and filtering phases, kGWASflow generates summary statistics for *k*-mer counts and visually displays the results through a variety of plots (Figure 1).

Next, kGWASflow generates the *k*-mer presence/absence genotype matrix, using the filtered *k*-mers obtained from all individuals/samples and utilizing the “build_kmers_table” function from the kmersGWAS library. This binary *k*-mers presence/absence table features *k*-mers as rows and individuals as columns. kGWASflow also allows users to convert this binary *k*-mers table into PLINK (Chang *et al*. 2015) format. After creating the *k*-mer table, kGWASflow constructs a kinship-relatedness matrix based on either the *k*-mers table or a user-provided SNP file in PLINK format. Users can choose between these two options via the configuration file. If users opt for a k-mer-based kinship matrix, kGWASflow executes the “emma_kinship_kmers” function from the kmersGWAS library, generating an EMMA-based (Kang *et al*. 2008) relatedness matrix. Alternatively, if an SNP-based kinship matrix is preferred, the pipeline employs the “emma_kinship” function from the same library. Users can specify the minor allele frequency and minor allele count parameters for this step by modifying the config file.

Upon generating the *k*-mers table and the kinship matrix, kGWASflow proceeds to perform k-mer-based GWAS utilizing the methodology developed by Voichek et al. (2020). In this phase, the *k*-mers table, kinship matrix, and phenotype file serve as inputs. The kmersGWAS method is applied independently to each provided phenotype using the “kmers_gwas.py” script (Voichek and Weigel 2020). In short, this method initially permutes the given phenotype N times (with N specified by the user in kGWASflow) and employs a linear mixed model (LMM) to associate *k*-mer presence/absence patterns with the phenotype and its permutations. By utilizing this approximated model, the top-ranking *k*-mers (determined by the user in kGWASflow, with a default value of 10,000) are identified and subsequently passed to the next stage. Afterward, these top *k*-mers are utilized as input in the following step, where the actual model implemented in GEMMA (Zhou and Stephens 2012) is employed to generate exact p-values for the phenotype and its permutations. The kinship matrix is used to account for the relatedness between individuals. A permutation-based 5% family-wise error-rate threshold is identified, and the *k*-mers surpassing this threshold are deemed statistically significant. After the completion of the kmersGWAS run, kGWASflow produces a results summary table (Figure 1), which incorporates the significant *k*-mers derived from the kmersGWAS stage.

#### Post-GWAS analysis

Post-GWAS analyses are all optional and highly dependent on the user’s preference, the organism of interest, and the research question. Users can easily activate or deactivate various Post-GWAS options within the configuration file. This phase concentrates on identifying the genomic location and context of significantly associated *k*-mers. This step can offer deeper insights into the context of previously recognized associations, while also aiding in the discovery of new associations that may contain a broader range of genetic variants. kGWASflow includes 3 Post-GWAS analysis options, which differ based on whether the user wants to map the kmers themselves, the reads they originated from, or contigs assembled from those reads.

The first post-GWAS analysis option is to directly map the trait-associated *k*-mers to a reference genome FASTA file (as defined in the configuration file). To map significant *k*-mers to a reference genome, users have the option to select from two read-mapping algorithms: bowtie or bowtie2 (Langmead and Salzberg 2012). kGWASflow maps the associated *k*-mers to a given genome FASTA using the preferred alignment algorithm and then converts the results into a sorted and indexed BAM file using samtools (Li *et al*. 2009). Finally, kGWASflow generates a Manhattan plot by incorporating the p-values from the kmersGWAS step and the genomic locations of the aligned *k*-mers obtained during this mapping stage.

The second option is to identify which sequencing reads the significant *k*-mers came from and map those reads to the genome FASTA file. When this option is enabled, kGWASflow initially retrieves the source reads for each associated *k*-mer from the FASTQ files of samples containing those *k*-mers. kGWASflow executes the”fetch_source_reads.py” script, which incorporates the “fetch_reads_with_kmers” tool (https://github.com/voichek/fetch_reads_with_kmers) at its core to identify the source reads of each trait-associated *k*-mer. After finding the source reads of trait-associated *k*-mers, the pipeline first merges and then sorts the reads using seqkit (Shen et al. 2016). kGWASflow maps the sorted reads to a reference genome FASTA file using bowtie2 (Langmead and Salzberg 2012) with “--very-sensitive-local” parameters. Alignments are filtered based on a mapping quality score defined by the user. kGWASflow also converts the alignment outputs into BAM and BED files for down-stream analysis. Optionally, the workflow generates IGV reports of the alignment results in HTML format using the igv-reports tool (https://github.com/igvteam/igvreports).

The third option also utilizes the source reads of the associated *k*-mers. If not previously obtained, the source reads of *k*-mers are retrieved as outlined above. In this step, instead of mapping the raw reads to a reference genome, kGWASflow first performs a de-novo assembly of the source reads using SPADES (Prjibelski et al. 2020) with the “--careful” parameter. After the assembly step, kGWASflow runs minimap2 (Li 2018) to map assembled contigs onto a reference genome FASTA file. Users also have the option to perform a BLAST (Altschul *et al*. 1990) query using blastn on the resulting contigs. The output of this step is the sorted, indexed, and quality-filtered (user-defined) BAM files of mapped contigs. If the BLAST option is enabled, the workflow outputs a BLAST results file. As in the previous step, kGWASflow optionally generates IGV reports of the contig mapping results using the igv-reports software.

## Results and discussion

To illustrate our workflow, we selected two distinct datasets that had been previously analyzed using different k-mer-based GWAS methods. The first dataset is a public *Escherichia coli* (*E. coli*) ampicillin resistance dataset that contains 241 strains of *E. coli* (Earle *et al*. 2016), which was used by Rahman et al. (2018) to test their k-mers-based association mapping tool HAWK (Rahman *et al*. 2018). Among 241 strains, 189 had ampicillin resistance and 52 were susceptible. By applying our workflow to this dataset, kGWASflow yielded results comparable to the findings from Rahman et al. (2018) (Supplementary Data S1: Tables 1,2). Figure 3 shows examples of kGWASflow output from this test run, including *k*-mer count summary statistics (Figure 3A-C) and k-mers-based GWAS results (Figure 3D, E). The configuration files and the HTML summary report from this kGWASflow run can be found in the Supplementary Data (Data S6-10). This *E. coli* ampicillin resistance dataset is included with the pipeline as the test dataset, and its results can be reproduced by executing a single command:

**Figure 3.**
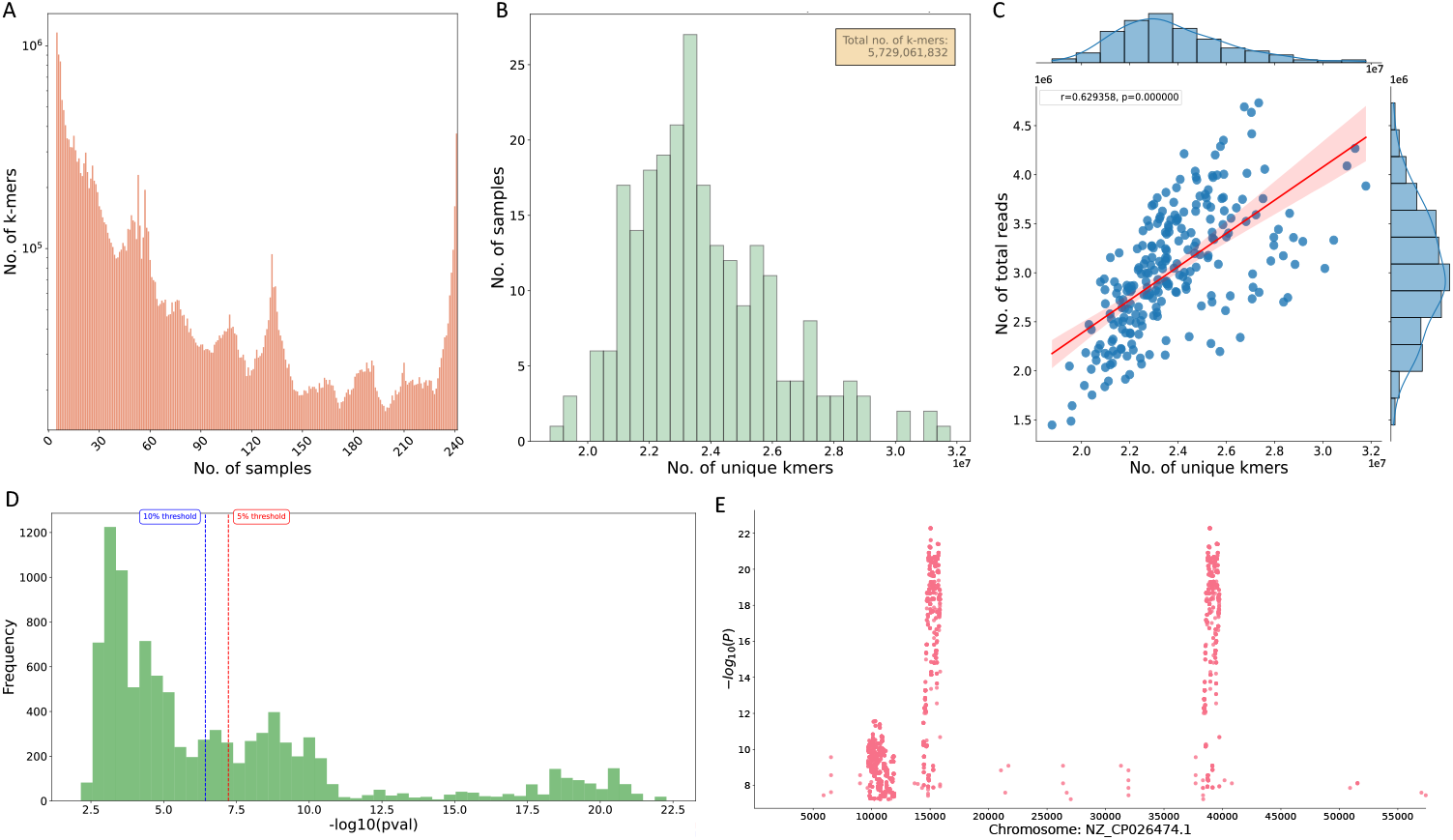
Example outputs obtained from kGWASflow by processing the *E. coli* ampicillin resistance dataset (Rahman et al. 2018). **(A)** Bar plot showing the number of *k* -mers that appeared in exactly ‘N’ number of samples (‘N’ goes between 1 to the total number of samples). Only the *k* -mers that passed the initial filtering step were used. **(B)** Histogram plot showing the distribution of non-canonical *k* -mer counts. The x-axis shows the unique *k* -mer counts and the y-axis shows the number of samples. The legend at the top right shows the total number of unique *k* -mers (non-canonical). The histogram plot for canonical counts can be found in the Supplementary Data (Figure S1A). **(C)** Joint plot showing the relationship between the non-canonical unique *k* -mer counts and the number of reads. The x-axis represents the number of unique *k* -mers (non-canonical), and the y-axis represents the number of total reads. The red line represents the linear regression line. The *r* -value is the Pearson correlation coefficient, and the *p*-value is the two-tailed *p*-value. The marginal distributions of the x and y axis are also shown on the top and right sides of the plot, respectively. The joint plot for canonical counts can be found in the Supplementary Data (Figure S1B). **(D)** Histogram of the -log10 *p*-values of each *k* -mer that passed the first kmersGWAS step. The red dashed line indicates the 5% family-wise errorrate threshold, while the blue dashed line indicates the 10% family-wise error-rate threshold. Only the *p*-values of the best *k* -mers from the first kmersGWAS step are used. *p*-values are obtained from GEMMA during the second step of kmersGWAS (a detailed explanation can be found in the k-mers based GWAS section). **(E)** Manhattan plot showing -log10 *p*-values of *k* -mers that are significantly associated with ampicillin resistance, mapped to their genomic locations. *k* -mers were mapped to *E. coli* plasmid pKBN10P04869A reference genome (PRJNA430286) using bowtie2.

~~~
    kgwasflow test --threads 16 \
                    --work-dir working_dir
~~~

Executing this command will trigger an automated download of the sequencing reads of all 241 strains of *E. coli* from NCBI, followed by QC analysis and the generation of a MultiQC report. Upon completion of the QC step, KGWASflow will proceed to the *k*-mer counting phase, subsequently executing *k*-mer filtration, *k*-mer table construction, and the generation of a kinship matrix based on *k*-mers. The workflow will then perform kmersGWAS using the k-mers table and the kinship matrix. kGWASflow will subsequently produce a summary of the kmersGWAS results, then execute all three post-GWAS phase options (Figure 1), successfully completing the test run. Ultimately, to produce an HTML report summarizing the workflow and the results, users only need to execute a single command after the test workflow run is completed:

~~~
    kgwasflow test --threads 16 \
                   --work-dir working_dir \
                   --generate-report
~~~

We employed a second dataset consisting of whole genome sequencing data of 261 maize lines from the Goodman-Buckler Maize Association Panel (Flint-Garcia *et al*. 2005) and three different maize phenotypes, including kernel color, upper leaf angle, and cob color. He et al. (2021) previously used this dataset to test their k-mer-based GWAS tool, which relies on *k*-mer occurrence count (KOC) rather than presence/absence (He *et al*. 2021). As in the previous test run, using kGWASflow on this dataset generated results that closely mirrored the findings of He et al. (2021). We identified trait-associated *k*-mers that passed the p-value threshold for each of the three phenotypes (Supplementary Data S1: Tables S3-8). Furthermore, using the post-GWAS analysis module of kG-WASflow, we have determined the probable genomic locations of these trait-associated *k*-mers and the results were comparable to He et al. (2021) (Supplementary Data S2: Figures S2-5). The configuration files and the HTML report from this kGWASflow run can be found in the Supplementary Data (Data S11-15).

## Conclusion

kGWASflow is an easy-to-install, reproducible, scalable, and user-friendly workflow written in Snakemake. It employs the kmersG-WAS method (Voichek and Weigel 2020) to conduct k-mer-based GWAS while offering enhanced pre- and post-GWAS analysis capabilities. kGWASflow offers extensive customization, either via the command line or a configuration file, enabling users to modify the workflow to their specific requirements. It takes advantage of Snakemake’s parallelization and scalability capabilities, making the workflow deployable in local computers, high-performance computing, or cloud computing environments. By utilizing Conda, kGWASflow effectively circumvents software dependency issues and prevents library/version conflicts. The easy expansibility of kGWASflow enables seamless integration of future enhancements and the incorporation of new or improved k-mers-based GWAS methods. Taken together, by creating a modular, customizable, and user-friendly workflow, we aim to enhance the accessibility and streamline k-mer-based GWAS, empowering a broader research community to leverage this approach.

## Data availability

kGWASflow is freely available (under MIT license) from GitHub (https://github.com/akcorut/kGWASflow) and also on Bioconda (https://anaconda.org/bioconda/kgwasflow), including necessary inputs to perform a test run. The GitHub repository contains a wiki (https://github.com/akcorut/kGWASflow/wiki) explaining in detail how to install, configure, and use the workflow.

## Supplemental data

**Supplementary Data S1:** Supplemental information contains Tables S1-S8.

**Supplementary Data S2:** Supplemental information contains Figures S1-S5.

**Supplementary Data S3-5:** Example default configuration files of kGWASflow.

**Supplementary Data S6-10:** The configuration files, phenotype information and HTML summary report of kGWASflow run with the E.coli dataset.

**Supplementary Data S11-15:** The configuration files, phenotype information and HTML summary report of kGWASflow run with the maize dataset.

Supplemental material available at FigShare: **insert_url_here**

## Acknowledgments

The majority of the computation for developing and testing the pipeline was conducted on the high-performance computing (HPC) resource SAPELO2 at the Georgia Advanced Computing Resource Center (GACRC).

